# Biofilm development on urinary catheters promotes the appearance of viable but non-culturable (VBNC) bacteria

**DOI:** 10.1101/2020.12.23.424279

**Authors:** Sandra A. Wilks, Verena V. Koerfer, Jacqui A. Prieto, Mandy Fader, C. William Keevil

**Author notes:** Address correspondence to: Sandra A. Wilks. Sandra A Wilks designed the study, carried out experimental work and prepared the manuscript. Verena V. Koerfer carried out experimental work and data analysis. Jacqui A. Prieto & Mandy Fader provided clinical expertise and reviewed content. C. William Keevil designed the study and reviewed the manuscript.

## Abstract

Catheter-associated urinary tract infections have serious consequences, both for patients and in impacting on healthcare resources. Much work has been carried out to develop an antimicrobial catheter. Although such developments have shown promise under laboratory conditions, none have demonstrated a clear advantage in clinical trials.

Using a range of microbiological and advanced microscopy techniques, a detailed laboratory study comparing biofilm development on silicone, hydrogel latex and silver alloy coated hydrogel latex catheters was carried out. Biofilm development by *Escherichia coli, Pseudomonas aeruginosa* and *Proteus mirabilis* on three commercially available catheters was tracked over time. Samples were examined with episcopic differential interference contrast (EDIC) microscopy, culture analysis and staining techniques to quantify viable but non-culturable (VBNC) bacteria.

Both qualitative and quantitative assessment found biofilms to develop rapidly on all three materials. EDIC microscopy revealed the rough surface topography of the materials. Differences between culture counts and quantification of total and dead cells demonstrated the presence of VBNC populations, where bacteria retain viability but are not metabolically active.

The use of non-culture based techniques showed the development of widespread VBNC populations. These VBNC populations were more evident on silver alloy coated hydrogel latex catheters, indicating a bacteriostatic effect at best. The laboratory tests reported here, that detect VBNC bacteria, allow more rigorous assessment of antimicrobial catheters offering an explanation for why there is often minimal benefit to patients.

**IMPORTANCE:** Several antimicrobial urinary catheter materials have been developed but, although laboratory studies may show a benefit, none have significantly improved clinical outcomes. The use of poorly designed laboratory testing and lack of consideration to the impact of VBNC populations may be responsible. While the presence of VBNC populations is becoming more widely reported, there remains a lack of understanding of the clinical impact or influence of exposure to antimicrobial products. This is the first study to investigate the impact of antimicrobial surface materials and the appearance of VBNC populations. This demonstrates how improved testing is needed prior to clinical trials uptake.

## INTRODUCTION

Urinary tract infections (UTIs) are the second most frequent cause of healthcare associated infections among hospitalized patients across Europe, with 60% attributable to indwelling urinary catheterization (catheter associated UTI, CAUTI). The use of a catheter increases the likelihood of bacteriuria [1,2]. Indeed, recent microbiome research in the healthy bladder [3,4] has shown urine not to be sterile, with asymptomatic bacteriuria routinely found when advanced molecular sequencing is used. Bacteriuria and the presence of a catheter result in a high risk of biofilm development, where bacteria attach to the catheter material, forming complex communities and increasing antibiotic resistance. Biofilms are known to have a role in CAUTI development and in catheter blockage, commonly caused by the presence of urease-producing bacteria.

The role of biofilms has been considered in laboratory-based studies [5,6,7,8,9,10,11] and there has been considerable work to develop antimicrobial materials [12,13,14,15]. While several have shown promise during laboratory testing, few have been assessed clinically. An exception is the use of silver alloy coated/impregnated catheters which became common in clinical settings, with *in vitro* evidence indicating a reduction in the incidence of CAUTI [16,17,18]. However, in a large-scale clinical trial involving approximately 7000 patients, Pickard et al. [19] demonstrated no difference in the incidence of infection between standard, uncoated catheters and silver impregnated/coated catheters in patients undergoing short-term catheterization. The trial did, however, note a reduction in bacteriuria in patients using the silver catheters.

This raises important questions as to why the trial data differed from what laboratory studies predicted. It would be expected that a polymicrobial community and the presence of human tissue *in vivo* would affect activity compared to controlled laboratory conditions. Also, it is known that bacteriuria does not inevitably lead to infection. However, the analytical techniques used and metabolic state of the bacteria can also have an impact and these are often neglected in studies.

To assess the antimicrobial activity of a material, attached cells are often removed and placed on nutritious agar media. This can lead to an underestimation of a biofilm population due to inefficiencies in removal and the presence of viable but non-culturable (VBNC) bacteria. VBNC bacteria arise from cells being sub-lethally stressed and being unable to grow on rich nutrient media, thus leading to an underestimation of population density [20,21,22]. VBNC bacteria can retain infectivity [22,23] and may be implicated in chronic, recurring infections as this metabolic state can be induced by the action of antibiotics [21,24]. Previously described methods to study biofilm development on urinary catheters can be limiting, as outlined recently [11]. It is also important to consider that bacteria within a biofilm state can have an altered phenotype and behave differently than their planktonic counterparts.

In the current study, a combination of *in situ* advanced microscopy [11,23] and *ex situ* viability staining and culture analyses, have been used to track biofilm development by three commonly found and clinically important bacterial uropathogens (*Escherichia coli, Pseudomonas aeruginosa* and *Proteus mirabilis*), on three catheter materials: silicone, hydrogel latex and silver alloy coated hydrogel latex, in a physiologically correct artificial urine medium. Using techniques specific to the detection of biofilms and VBNC bacteria, we illustrate how different catheter materials affect bacterial attachment, biofilm formation and can lead to increased VBNC populations.

## RESULTS

### Unused catheters

EDIC microscopy allowed rapid examination of silicone, hydrogel latex and silver alloy coated catheters (Figure 1a-c). The silicone catheter (Figure 1a) had a smooth topography but there was evidence of pitting and undulations as well as parallel striations formed by extrusion during the manufacturing process. In contrast, the hydrogel and silver alloy coated catheters (Figures 1b & c) had highly disordered, rough surface topographies. For all three materials, there were numerous potential bacterial attachment sites.

**Figure 1.**
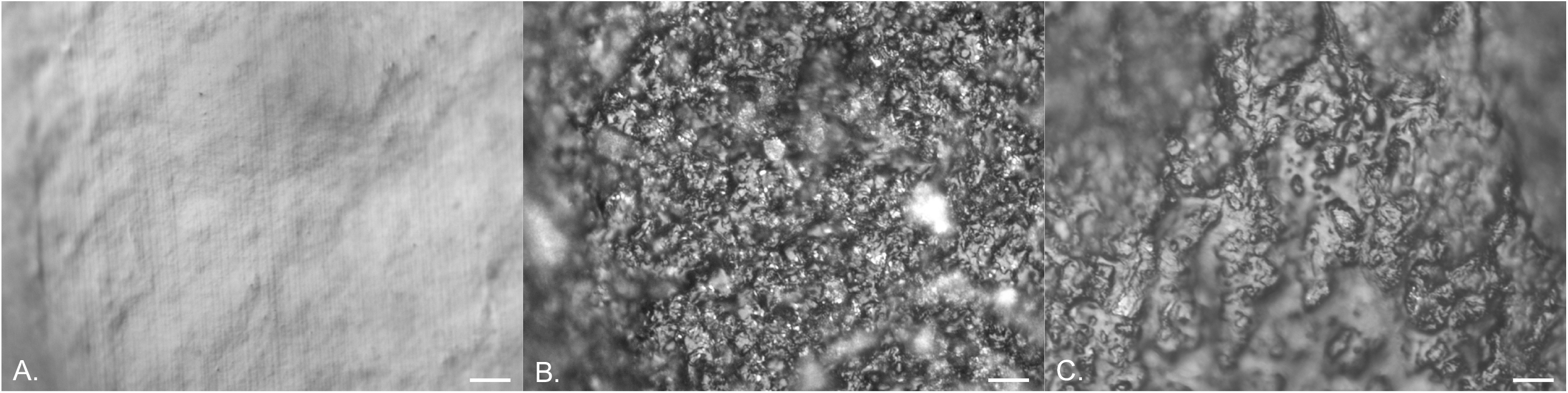
EDIC images of the surfaces of clean, unused catheters. a. silicone catheter. b. hydrogel latex catheter. c. silver alloy coated hydrogel latex catheter. (Magnification x 1000, bar = 10 μm).

### Qualitative assessment of biofilm development

#### Silicone

It was possible to use EDIC microscopy to visualize and track biofilm development on silicone catheters. Figures 2a-f show an example sequence from initial attachment progressing to biofilm development when *E. coli* was inoculated into artificial urine. Over the first 6 h exposure, there was a gradual increase in attached bacteria and material, from individual cells (2 h) to clear clustering and increasing density by 6 h (Figures 2a-c). Following 6 h exposure, distinct microcolony development was seen with associated extracellular polymeric substances (EPS) forming. After 24 h, a thick layer covered almost the entire surface and this continued over 48 h (Figures 2d & e). When sections were left for 72 h, channels and areas where biofilm had sloughed away were apparent, suggesting nutrient limitation, detachment events and exposed areas ready for re-colonisation (Figure 2f).

**Figure 2.**
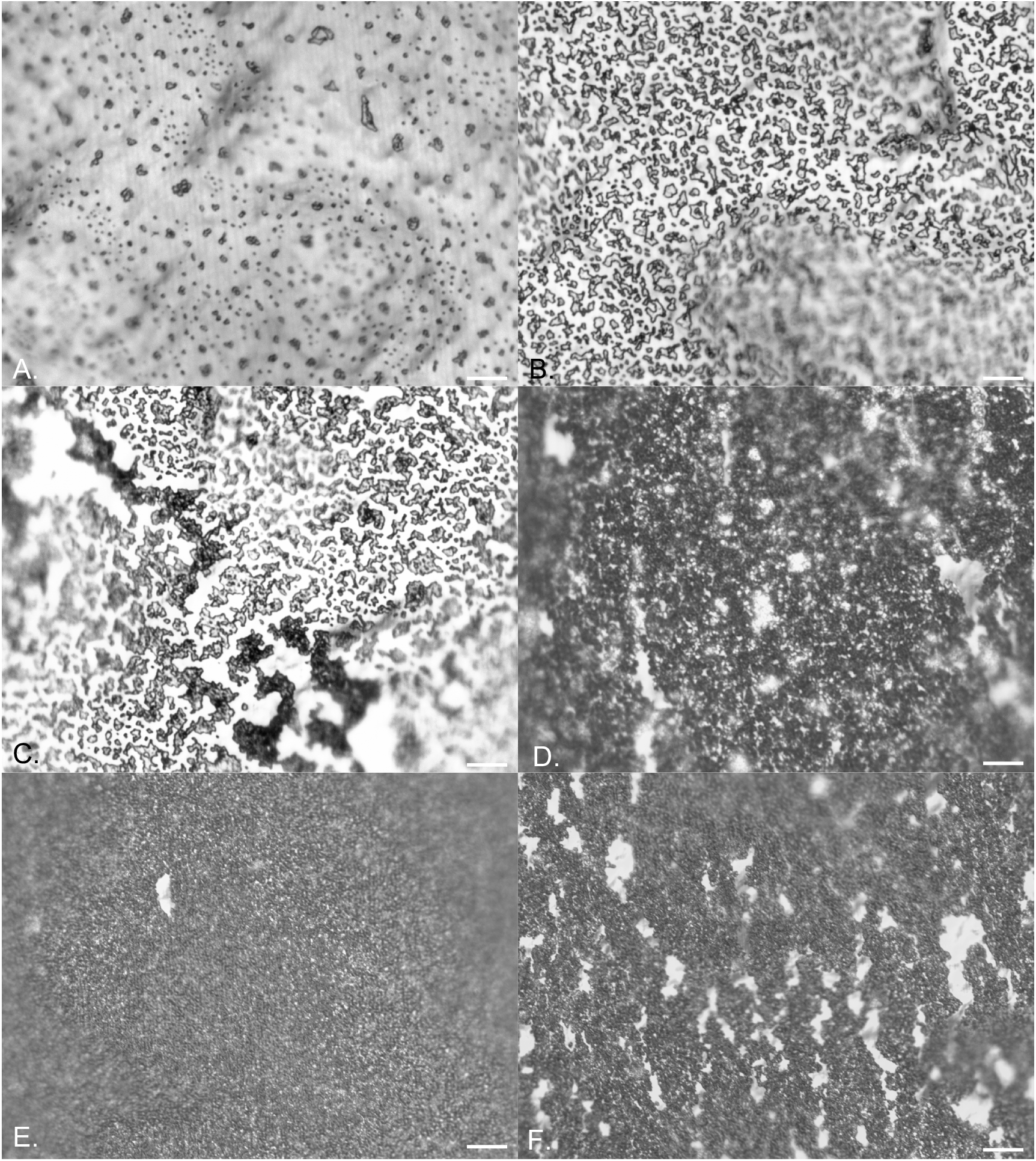
EDIC images showing attachment of *E. coli* to silicone catheters in artificial urine. a. 2 h, b. 4 h, c. 6 h, d. 24 h, e. 48 h, f. 72 h exposure. (Magnification x 1000, bar = 10 μm).

Similar results were obtained for both *P. aeruginosa* (Figures 3a-f) and *P. mirabilis* (Figures 4a-f). *P. aeruginosa* rapidly colonised the surface, leading to increased microcolony formation and large amounts of extra polymeric substances (EPS). After 24 h exposure, thickening of the biofilm was observed with stacks of microcolonies extending out from the surface (Figure 3d). There was no evidence of sloughing or detachment at increased exposure times (Figures 3e & f) but areas with increased reflectance indicated crystal formation and deposition due to urease activity. For *P. mirabilis*, single cell attachment occurred rapidly and after 6 h exposure an open mosaic structure was observed (Figure 4c). As reported previously [11], this was followed by almost complete coverage of the surface with evidence of crystal formation clear by 48 h and after 72 h, large areas had sloughed away, creating sites for re-colonisation (Figures 4d-f).

**Figure 3.**
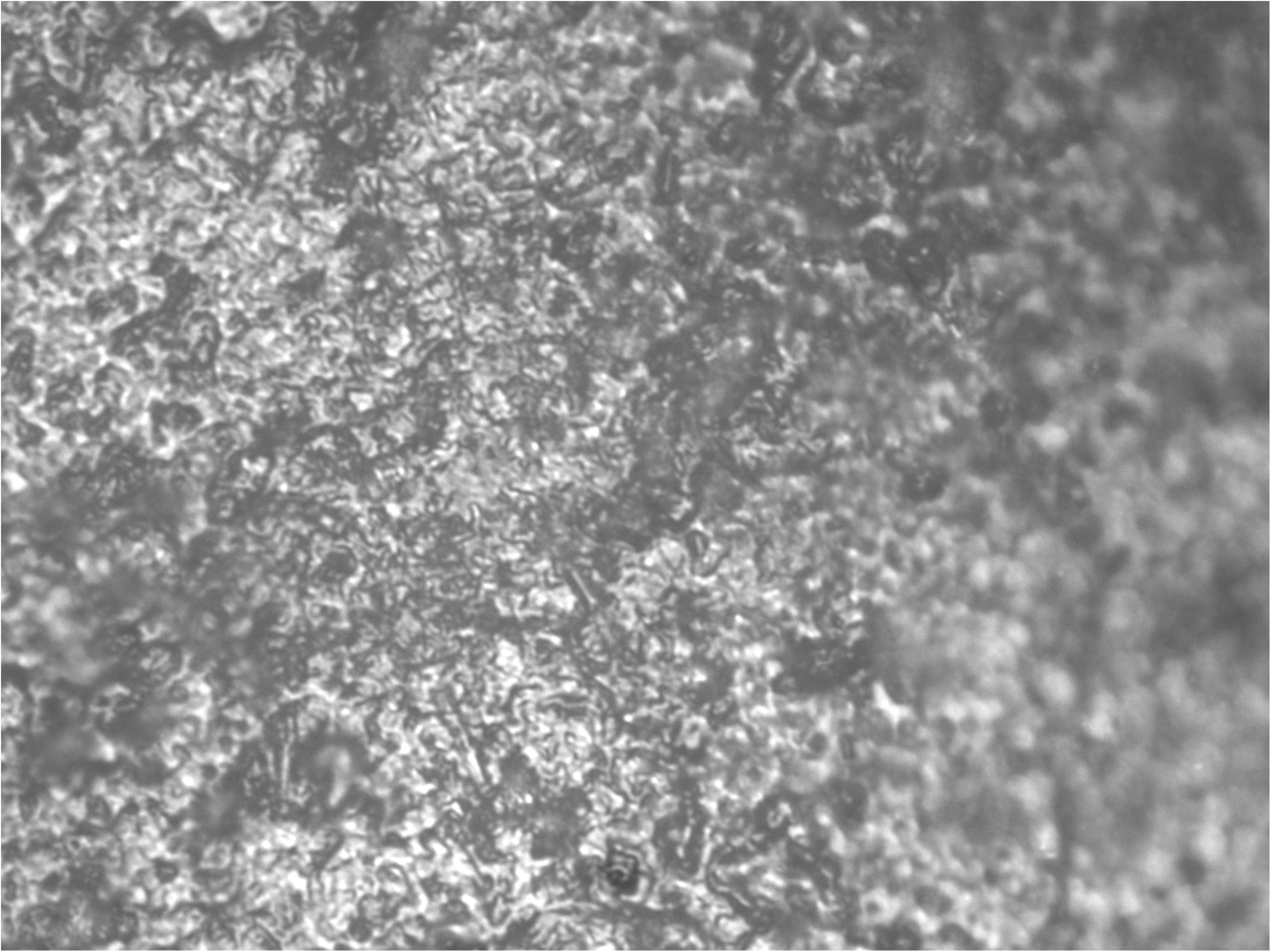
EDIC images showing attachment of *P. aeruginosa* to silicone catheters in artificial urine. a. 2 h, b. 4 h, c. 6 h, d. 24 h, e. 48 h, f. 72 h exposure. (Magnification x 1000, bar = 10 μm).

**Figure 4.**
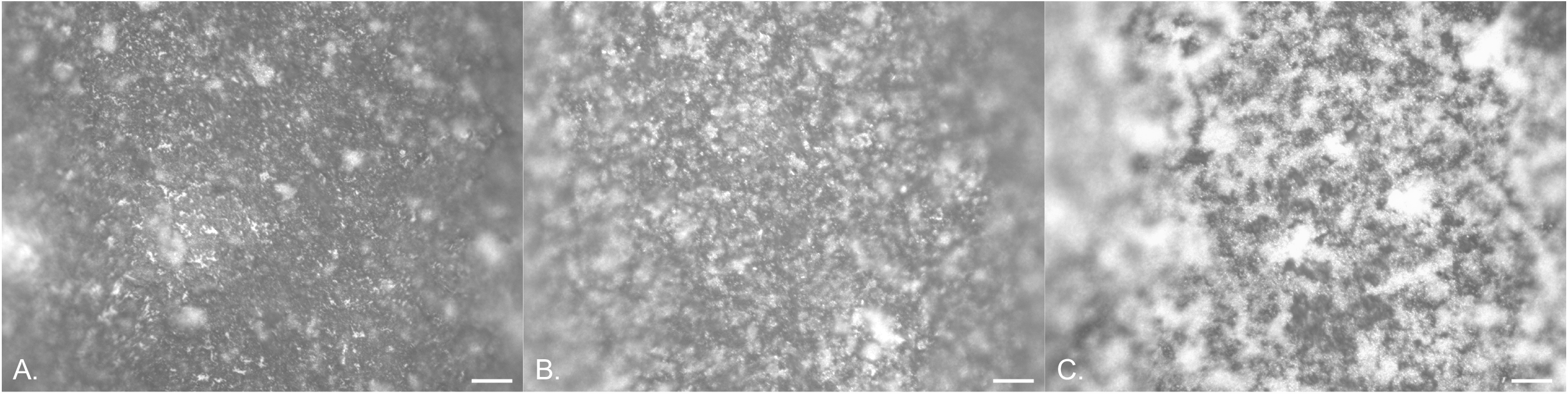
EDIC images showing attachment of *P. mirabilis* to silicone catheters in artificial urine. a. 2 h, b. 4 h, c. 6 h, d. 24 h, e. 48 h, f. 72 h exposure. (Magnification x 1000, bar = 10 μm).

#### Hydrogel latex/silver

It was not possible to follow the early stages of bacterial attachment on either material due to the highly disordered surface topographies. Some evidence of crystalline biofilm development could be seen over longer exposure times (24, 48 and 72 h exposure) for *P. aeruginosa* (Figures 5a-c) and *P. mirabilis* (Figures 6a-c) due to urease production.

**Figure 5.**
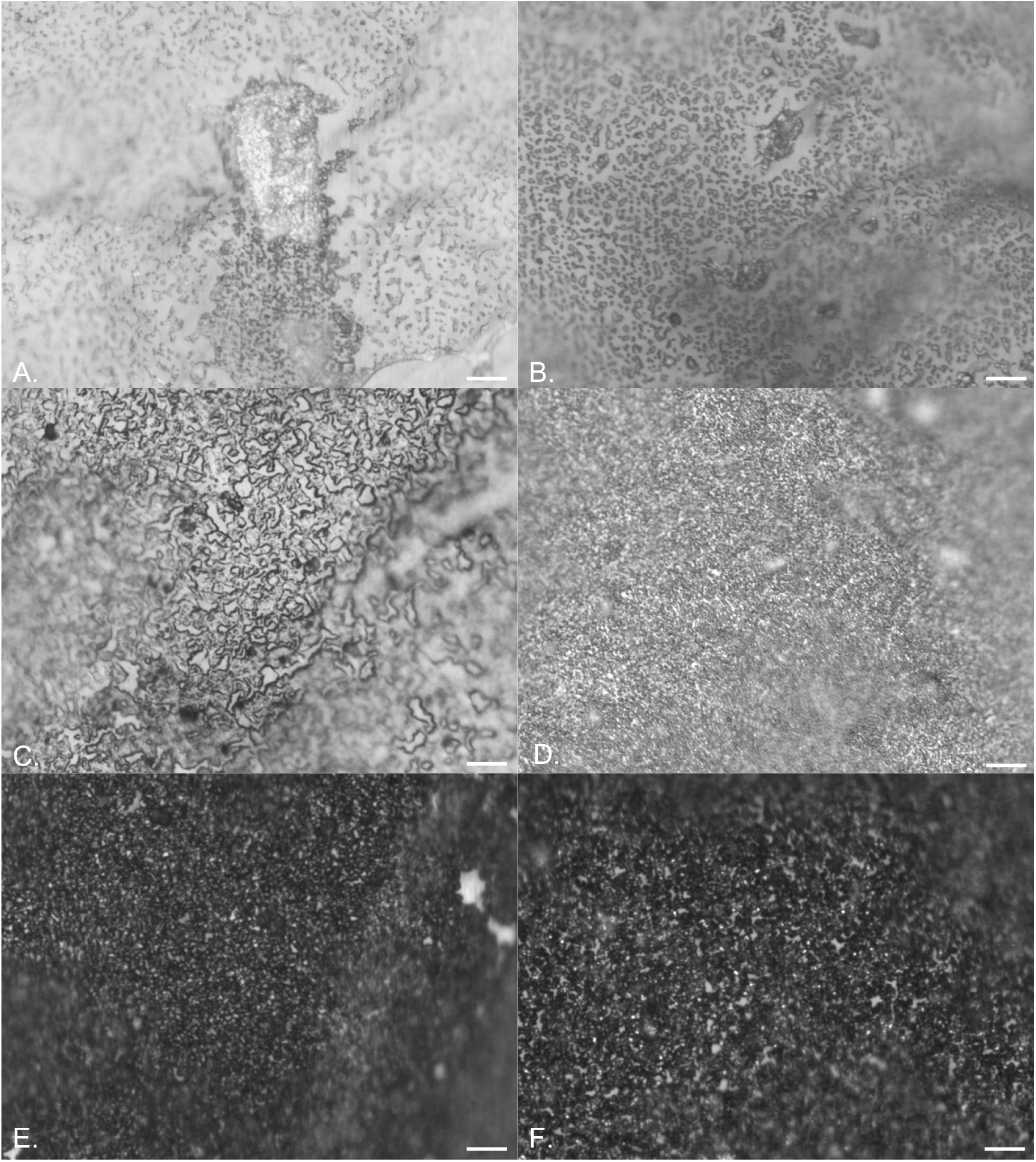
EDIC images showing attachment of *P. aeruginosa* to hydrogel latex catheters in artificial urine. a. 24 h, b. 48 h, c. 72 h exposure. (Magnification x 1000, bar = 10 μm).

**Figure 6.**
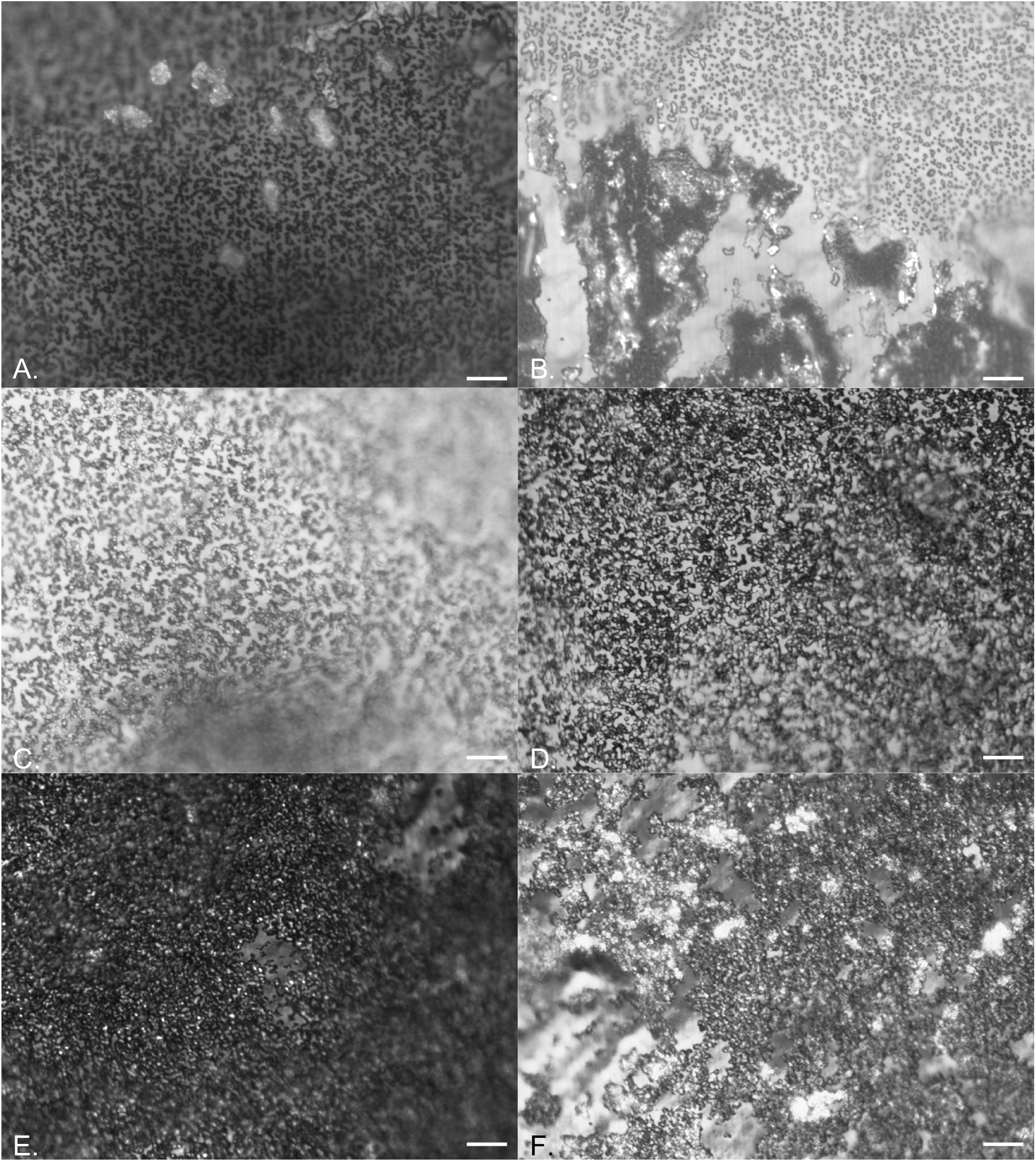
EDIC images showing attachment of *P. mirabilis* to silver alloy coated hydrogel latex catheters in artificial urine. a. 24 h, b. 48 h, c. 72 h exposure. (Magnification x 1000, bar = 10 μm).

### Quantitative assessment of biofilm formation

Three methods were used to quantify biofilm development: staining with SYTO 9 (total cell counts, TCC), staining with PI (dead cells, dead) and culture analysis (colony forming units, cfu).

### E. coli

The attachment and development of *E. coli* biofilms over 72 h was assessed and quantified with results shown in Figures 7a-c, for silicone, hydrogel latex and silver alloy coated hydrogel latex respectively, with the percentage of the population in a VBNC state given (refer to equation 1). There were no significant differences between cfu, TCC and dead counts over time or between catheter materials (*P* < 0.05).

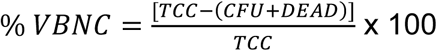

**Figure 7.**
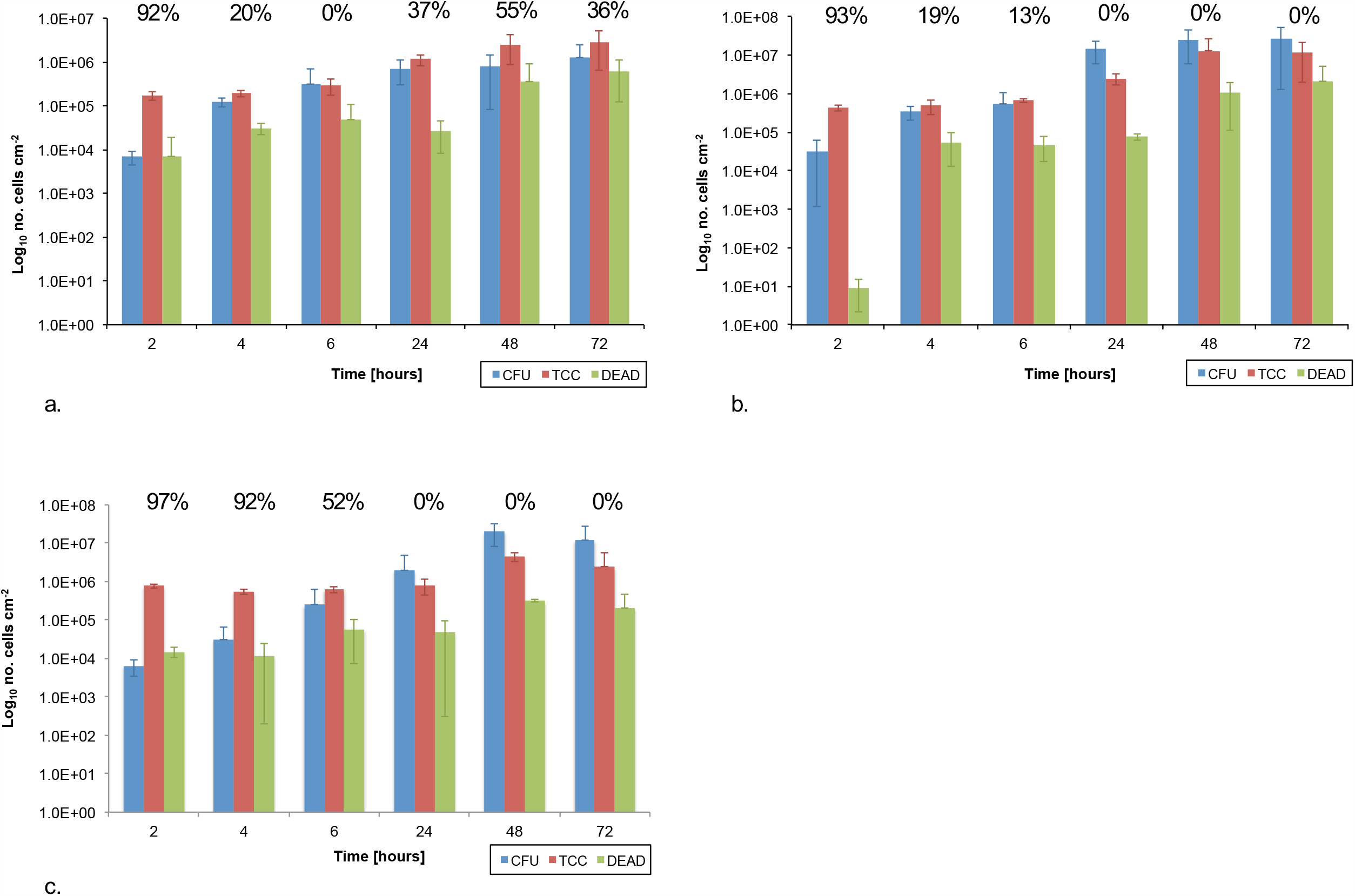
Graph showing the numbers of colony forming units (cfu), total cell counts (TCC) and dead cell counts (dead) per cm^2^ over time following exposure to *E. coli*. a. silicone b. hydrogel latex, c. silver alloy hydrogel latex catheters. The percentage of VBNC population (colony forming units plus dead cell counts) in relation to total cell count are shown.

Equation 1. Calculation of the percentage VBNC cells within total population where TCC = total number of cells (SYTO 9 labelled), CFU = colony forming units representing culturable cells, and DEAD = non-viable cells (PI labelled).

On silicone (Figure 7a), after 2 h exposure there was a greater than 1 log difference between the sum of cfu plus dead cells and the TCC counts, suggesting at least 90% of the population were VBNC. This was also observed after 48 and 72 h exposure, where there was an approximate 0.5 log difference. On hydrogel latex (Figure 7b), after 24 h and greater exposure times, the cfu counts were equal or greater than the TCC with many cells undergoing cell division when examined under the microscope. The early time points did show evidence of VBNC populations but these decreased from 93% after 2 h to 13% at 6 h. A different pattern was found on silver alloy coated hydrogel latex catheters (Figure 7c) where, although cfu counts ≥ TCC at 24 h and longer exposure time points, in the first 4 h the sum of cfu plus dead cells remained between 1 - 2 log lower than TCC, suggesting the presence of a VBNC population of > 90%, only decreasing to 52% at 6 h.

### P. aeruginosa

The attachment and development of *P. aeruginosa* biofilms over 72 h was assessed and quantified, with results shown in Figures 8a-c for silicone, hydrogel latex and silver alloy coated hydrogel latex respectively, with the percentage of the population in a VBNC state given (refer to equation 1). When comparing the three materials there were some significant differences (*P* < 0.05). Dead cell counts were significantly different at the longer exposure times ≥ 24 h, with silicone < silver alloy < hydrogel latex. However, at 48 h, TCC were also significantly different in the same order, with the highest values recorded for hydrogel latex. The only significant difference for cfu values was after 72 h where results on hydrogel latex (Figure 8b) were almost 2 log higher than for silver alloy coated hydrogel latex (Figure 8c).

**Figure 8.**
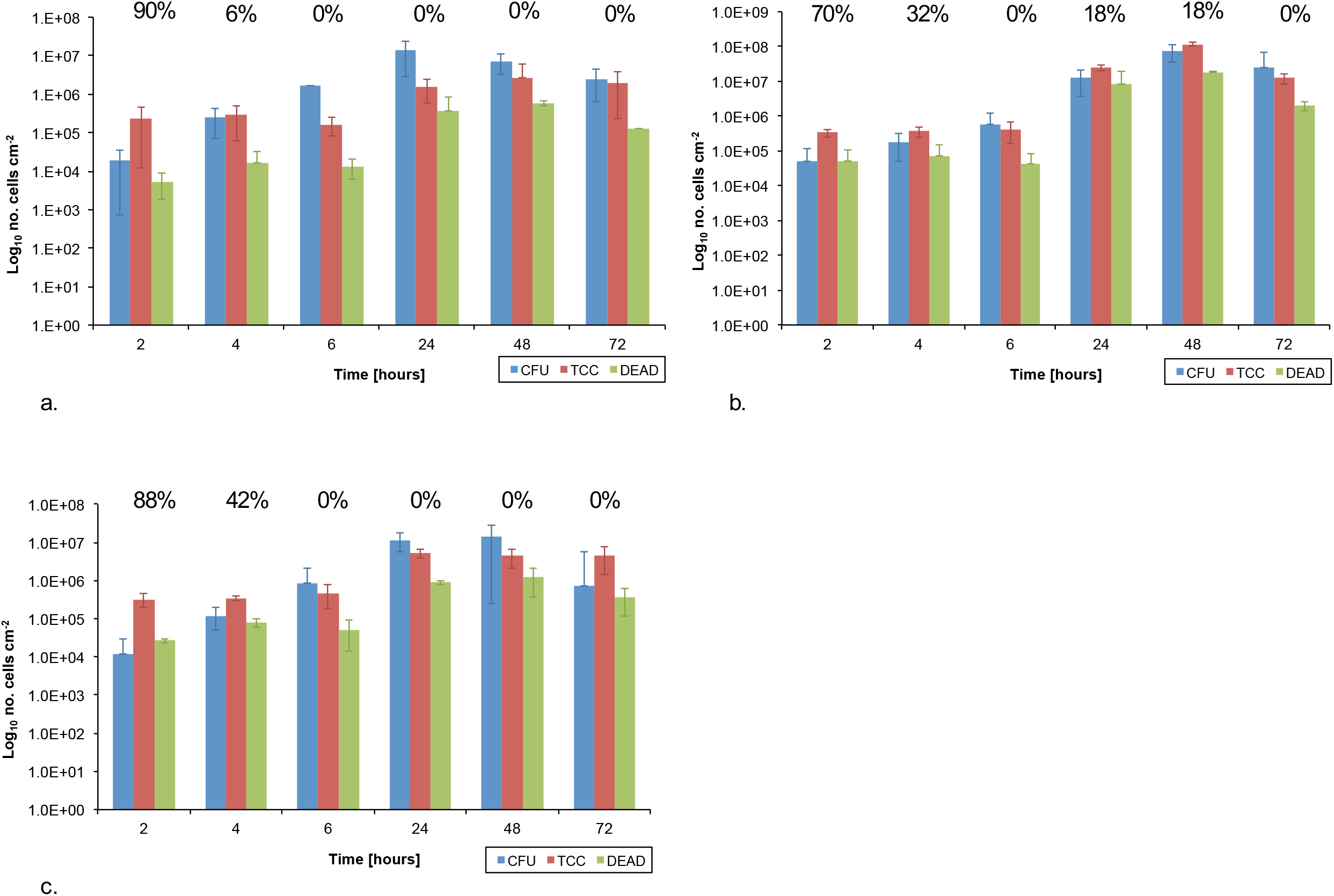
Graph showing the numbers of colony forming units (cfu), total cell counts (TCC) and dead cell counts (dead) per cm^2^ over time following exposure to *P. aeruginosa*. a. silicone b. hydrogel latex, c. silver alloy hydrogel latex catheters. The percentage of VBNC population (colony forming units plus dead cell counts) in relation to total cell count are shown.

On silicone (Figure 8a), at 2 h exposure, cfu values plus dead counts were over 1 log lower than TCC indicating the presence of VBNC cells (90% of the population). However, at longer time points, cfu counts were greater than TCC with many cells in the process of dividing and hence underestimated, with no evidence of VBNC formation. For hydrogel latex (Figure 8b), there was little difference between TCC and cfu, although 70% of the population were in a VBNC state at 2 h and 32% after 4 h. This pattern was followed but increased on silver alloy coated hydrogel latex (Figure 8c), cfu plus dead cells were approximately 1 log lower than TCC after 2 h with 88% of the population in a VBNC state reducing to 42% at 4 h.

### P. mirabilis

The attachment and development of *P. mirabilis* biofilms over 72 h was assessed and quantified, with results shown in Figures 9a-c for silicone, hydrogel latex and silver alloy coated hydrogel latex respectively, with the percentage of the population in a VBNC state given (refer to equation 1). When comparing materials, the TCC values were significantly different at all time points with silver alloy< silicone < hydrogel latex. The cfu and dead cell counts did not differ significantly across materials.

**Figure 9.**
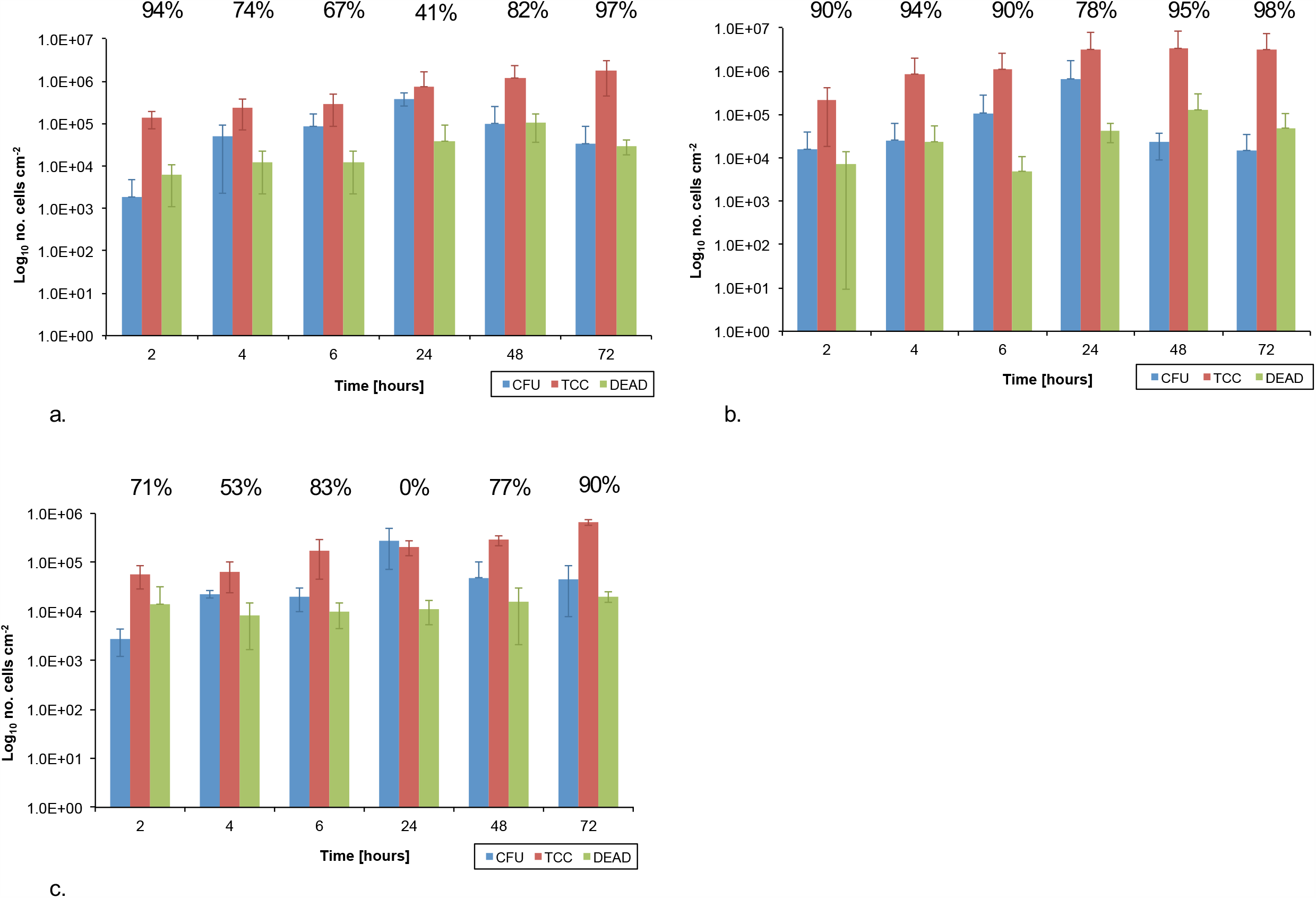
Graph showing the numbers of colony forming units (cfu), total cell counts (TCC) and dead cell counts (dead) per cm^2^ over time following exposure to *P. mirabilis*. a. silicone b. hydrogel latex, c. silver alloy hydrogel latex catheters. The percentage of VBNC population (colony forming units plus dead cell counts) in relation to total cell count are shown.

On silicone (Figure 9a), there were no significant differences over time; however, there was evidence of VBNC populations at all time points with an approximate 2 log difference between the sum of cfu plus dead cells and TCC after 2 and 72 h exposure times (94 and 97% respectively). This was also seen on hydrogel latex (Figure 9b) with an approximately 2 log (≤ 95%) difference at the longer exposure times of 48 and 72 h. Likewise, with the exception of 24 h, the number of cfu was always over 1 log (ranging between 53 - 90%) lower than the TCC on silver alloy coated hydrogel latex (Figure 9c).

## DISCUSSION

Despite continued efforts to produce effective antimicrobial catheter materials and coatings that resist biofilm development, the problems of CAUTI and blockage prevail. To date, no material has been found to improve clinical outcomes significantly over long-term use and many are described as being bacteriostatic rather than bactericidal.

The use of silver alloy coatings and impregnated catheters has been shown to reduce bacterial numbers in *in vitro* studies [16,17,18,25]. Indeed, Gabriel et al. [25] reported on the efficacy of a silver-coated catheter in 1996, leading to subsequent clinical approval. While silver-coated/impregnated catheters became widely adopted, several *in vivo* studies have indicated that any positive benefit of using silver coated or impregnated catheters is short-term only (< 7 days) [26,27,28,29], with a delay in colonization observed. In these cases, the antimicrobial action of silver can, at best, be described as bacteriostatic. This was also demonstrated in a large, randomised, multicentre trial [19] where no reduction in CAUTI following the use of silver alloy coated catheters compared to nitrofural-impregnated and PTFE-coated catheters was found. Results from this study indicated that to prevent one incidence of CAUTI at least 1000 people would need to be using the silver alloy coated catheter. This contradicts the findings of previous studies [30,31], which had led to the recommendation of silver alloy coated catheters for routine short-term use in the UK and USA. This questions whether appropriate *in vitro* studies could have prevented the subsequent recommendation for clinical use. Indeed, early studies such as by Johnson et al. [16] did report that alternative antimicrobial catheter materials, such as nitrofurazone containing, significantly outperformed silver hydrogel catheters.

In the current study we have tracked biofilm formation and investigated whether metabolically inactive, VBNC populations arise on three catheter materials, including a silver-alloy, and could be responsible for clinical outcome failures.

The VBNC state remains poorly understood although it has been reported to occur in a wide range of bacterial species covering most phyla [21]. The majority of work has focused on non-clinical environments including water [32,33,34] and food production [22], with several referencing biofilms as providing a reservoir niche. The impact of VBNC bacteria in clinical infection risk remains poorly understood with the role of persisters (cells that demonstrate antibiotic tolerance with restored growth on/in nutrient media once the stress is removed) more frequently studied [35]. There remains debate on the similarities and differences on the metabolic state of persister and VBNC cells [21,36,37], with a continuum in metabolic activity from dead to actively growing seeming likely [21]. Moreover, several studies [22,34] have demonstrated how VBNC cells, even in a ‘deeper’ level of dormancy can lead to infection in animal models. Additionally, it is well known that any medical device which enters the body provides a high-risk interface for bacterial attachment and biofilm formation. It is also known that both VBNC and persister cells can be commonly isolated from biofilms where stressors such as nutrient depletion, redox gradients and pH/ionic changes can occur.

Considering these factors, and our recent understanding of the diverse urinary microbiome [3], the potential for VBNC populations to form on urinary catheters is high, however this has not been explored previously. As CAUTI (particularly in chronic, recurring infections and blockages) and rapid biofilm development on urinary catheters are well-documented but no antimicrobial material (whether impregnated or coated) has been successful clinically in long-term patients, the appearance of VBNC populations may be an important factor. Indeed, the implications of VBNC populations in any device-related contamination and infection has not been widely considered or studied.

Using a combination of qualitative and quantitative methods, we tracked bacterial attachment, biofilm formation and the appearance of VBNC populations over time on three different catheter materials; silicone, hydrogel latex, and silver alloy impregnated hydrogel latex. The use of EDIC microscopy (11,38) demonstrated how, for all three species of bacteria tested, initial attachment was rapid, occurring in less than 2 h. *E. coli* showed biofilm maturation over the first 48 h before exhibiting a typical mosaic pattern with detachment events. In contrast, the biofilm-forming *P. aeruginosa* showed no signs of detachment, forming an extensive and thick biofilm with evidence of stack formation. The urease-producing species (*P. mirabilis* and *P. aeruginosa*) showed evidence of microcrystalline formation and in the former, to the development of complex crystalline encrustations, which lead to catheter blockages. This corresponds to work by Wilks et al. [11] where four distinct stages were identified.

Such qualitative observations are useful in understanding the susceptibility of catheter materials for biofilm development and gross structural characteristics but do not reflect differences in bacterial numbers or variations in viability and metabolic state. By using three quantitative methods; culture analysis (culturable cell count) and separate enumeration of SYTO 9 labelled (total cell count) and PI labelled (dead cell count) bacteria, we have shown the presence of an increasing VBNC population on silver alloy coated catheters. If no VBNC population is present, culture data plus the numbers of dead cells (PI labelled) should equal the total cell count (SYTO 9 labelled). While results for *P. aeruginosa* did follow this pattern on all materials (other than at short time points), indicative of its behavior as a strong biofilm-forming species, uropathogenic *E. coli* and *P. mirabilis* did not. This implies that VBNC cells are a natural component of the urinary catheter-related biofilms for these two bacteria. However, increased numbers of VBNC cells were observed on the silver alloy coated catheters and with little to no evidence of antimicrobial killing from this material type. Interestingly, a study by Zandri et al. [39] detected a VBNC population of *Staphylococcus aureus* in biofilms on central venous catheters removed from patients after implantation times of 3 days to 3 months (77% of 44 samples showed the presence of VBNC *S. aureus*).

The question arises why previous *in vitro* studies showed significant antimicrobial properties of silver alloy coated/impregnated catheters, which led to rapid clinical adoption. Consideration must be given to how experiments were designed and the measurements collected. These studies have relied on the growth of bacteria in nutritious laboratory media (where VBNC populations will be missed) or in minimal media; neither accurately reflecting the physiological environment of the urinary system. Studies have used radiolabelled leucine [17,25] to track bacterial adherence, and agar diffusion where inhibition to sections of catheter materials were measured [16,17] on Mueller-Hinton agar plates. Samuel and Guggenbichler [18] described four methods including growth of bacteria released from catheter surfaces by turbidity readings, measurement of antimicrobial activity by the Dow Shaker method (immersion in inoculated saline and aliquots of the suspension plated after set amounts of time), and the roll plate technique (inoculated catheter sections being rolled over agar plates). The current study utilises a physiologically correct artificial urine medium, which in itself influences the attachment and development of biofilms including the impact of urease release and crystal formation [11]. It then becomes clear how the mismatch between *in vitro* study design could have impacted on the discrepancies seen in *in vivo* use.

Although the clinical significance of VBNC bacteria is not fully understood, there is increasing evidence that they may have an important role in chronic and persistent infections (Li et al. 2014). The possibility of metabolically inactive bacteria having a role in UTI persistence was described by Mulvery et al. [40] who showed how a reservoir of inactive *E. coli* could be found inside bladder epithelial cells. The implications of such a population have not been explored further in relation to catheter-associated biofilm risk. Studies are demonstrating widespread retention of infectivity [22,34], indicating that the VBNC state could be a key mechanism and their increased antibiotic tolerance impacts on the efficacy of treatment plans. Indeed, the presence of a VBNC population, as demonstrated here, is a possible explanation for why silver alloy coated or impregnated catheters have failed in clinical trials. If these VBNC cells retain infectivity, this may account for why there was no reduction in CAUTI in the Pickard study [19], despite previous *in vivo* studies reporting a reduction in bacteriuria as found using standard urinalysis.

This work demonstrates the need for rigorous testing of medical device materials prior to clinical trial and market release, and full understanding of the implications of bacteriostatic versus bactericidal populations particularly considering the presence of VBNC populations. By combining several analysis methods, robust data can be obtained on the real antimicrobial activity of materials. This is vitally important in order to better understand the potential of new materials to reduce infection, blockage, improve patient care and quality of life, and reduce the financial burden of CAUTI.

## MATERIALS AND METHODS

### Bacterial inocula

*Escherichia coli* NCTC 9001, *Pseudomonas aeruginosa* PAO1 and *Proteus mirabilis* NCTC 10975 were grown overnight at 37°C in tryptone soya broth (TSB) (Oxoid, UK). Aliquots were centrifuged at 3780 x *g* for 10 min and pellets resuspended in artificial urine medium [41].

### Biofilm development on catheters

Three indwelling Foley catheters were tested: a 100% silicone (Rüsch), a hydrogel latex (Bard Biocath Hydrogel) and a silver alloy coated hydrogel latex (Bard Bardex® I.C.). These were cut longitudinally and transversely to give approximately 1 cm^2^ surface areas sections.

Catheter sections were placed in six-well tissue culture plates (two per well) and covered with 3 ml artificial urine [41]. To test wells, the inoculum was added (to give a final concentration of approximately 10^8^ cfu/ml), with each well representing a separate time point. Control samples did not have bacterial inocula added. These plates were incubated at 37°C. Samples were taken after 2, 4, 6, 24, 48 and 72 h exposure. Following removal from inoculated urine, each catheter section was gently washed with phosphate buffered saline (PBS).

### Qualitative assessment of biofilm development

One section was set aside for direct analysis by episcopic differential interference contrast (EDIC) microscopy using a Nikon Eclipse LV100D microscope with EXFO X-Cite 120 metal halide fluorescence system and long working metallurgical Nikon Plan Achromat objectives (Best Scientific, UK) [11,38].

### Quantitative assessment of biofilm development

The second catheter section from each well was used for indirect biofilm analysis. Biofilm was removed by scraping the surface with a sterile 1 µl inoculation loop which was transferred to 5 ml of PBS and vortex mixed for 30 sec. This resuspended biofilm was serially diluted in PBS, plated onto tryptone soya agar (TSA) (Oxoid, UK) and incubated at 37°C overnight. Resuspended biofilm samples were also stained with the SYTO9/ Propidium iodide (PI) LIVE/DEAD® BacLight™ system (Invitrogen, UK), to give total cell counts and numbers of dead cells. In each case, 1.5 µl of either SYTO 9 or propidium iodide (PI) was added to a 1 ml sample and incubated, in the dark, for 20 min. Following this, samples were filtered onto black 0.2 µm pore size diameter polycarbonate filters (Whatman, UK) and placed onto glass slides. Filters were examined, under epifluorescence illumination, using oil immersion and numbers of stained bacteria counted across a random selection of 10 fields of view.

### Statistical analysis

All experiments were repeated in triplicate. Results obtained for culturable bacteria, total and dead cell counts were log-transformed. Differences between analysis methods and catheter materials were assessed using a one-way analysis of variance (ANOVA) followed by Tukey’s multiple comparison test (Prism, GraphPad Software Inc.). Differences were considered significant if *P* < 0.05.

## ACKNOWLEDGEMENTS

This work was carried out by V. Koerfer as part of her Master’s degree from University of Duisberg-Essen and as part of an Institute for Life Sciences (University of Southampton) fellowship awarded to S.A. Wilks.

